# Timing of exposure is critical in a highly sensitive model of SARS-CoV-2 transmission

**DOI:** 10.1101/2021.12.08.471873

**Authors:** Ketaki Ganti, Lucas M. Ferreri, Chung-Young Lee, Camden R. Bair, Gabrielle K. Delima, Kate E. Holmes, Mehul S. Suthar, Anice C. Lowen

**Author notes:** These individuals contributed equally to the research presented (Ketaki Ganti, Lucas M. Ferreri, Chung-Young Lee, Camden R. Bair). These individuals contributed equally to the research presented (Gabrielle K. Delima, Kate E. Holmes).

## Abstract

Transmission efficiency is a critical factor determining the size of an outbreak of infectious disease. Indeed, the propensity of SARS-CoV-2 to transmit among humans precipitated and continues to sustain the COVID-19 pandemic. Nevertheless, the number of new cases among contacts is highly variable and underlying reasons for wide-ranging transmission outcomes remain unclear. Here, we evaluated viral spread in golden Syrian hamsters to define the impact of temporal and environmental conditions on the efficiency of SARS-CoV-2 transmission through the air. Our data show that exposure periods as brief as one hour are sufficient to support robust transmission. However, the timing after infection is critical for transmission success, with the highest frequency of transmission to contacts occurring at times of peak viral load in the donor animals. Relative humidity and temperature had no detectable impact on transmission when exposures were carried out with optimal timing. However, contrary to expectation, trends observed with sub-optimal exposure timing suggest improved transmission at high relative humidity or high temperature. In sum, among the conditions tested, our data reveal the timing of exposure to be the strongest determinant of SARS-CoV-2 transmission success and implicate viral load as an important driver of transmission.

## Introduction

The COVID-19 pandemic has led to a public health emergency and social disruption on a scale not seen since the influenza pandemic of 1918. Non-pharmaceutical interventions have been pursued in many parts of the world with the goal of limiting the impact of the outbreak (1-3). These interventions seek to interrupt transmission of the virus (1-3). Effective use of non-pharmaceutical interventions therefore relies heavily on fundamental understanding of SARS-CoV-2 transmission and the viral, host, and environmental factors that modulate its efficiency.

For example, efforts to contain viral spread through contract tracing rely on estimates of the duration of exposure needed to support transmission. In defining a close contact, exposure to an infected individual for a minimum of 15 or 30 minutes is often used (4, 5). In turn, quarantine measures for those identified as a close contacts rely on estimates of incubation period, commonly reported as 10–14 days (6, 7). Similarly, policies for isolation of positive cases reflect the potential for onward transmission within a period up to 10 or 14 days post-infection (6, 7). Evidence-based refinement of these definitions is important for establishing infection-control practices that minimize risk of transmission while also mitigating the social and economic burden of quarantine measures.

Much attention has also been paid to the potential for various environmental conditions to modulate the efficiency of transmission. Epidemiological investigation of environmental parameters has returned varied results (8-10). Low temperature was found to be associated with increased SARS-CoV-2 transmission in some cases (11, 12) but not others (13, 14). Similarly, dry conditions were found to be favorable for SARS-CoV-2 spread in a subset of studies (12, 15). In general, detected effects of temperature and humidity on reproduction number or epidemic growth were dwarfed by those of active interventions such as restrictions on mass gatherings (14, 16, 17). Indeed, the complexity of conditions under which human exposures occur, and incomplete information regarding those conditions, can frustrate efforts to define parameters important for transmission efficiency. Examination of transmission in relevant experimental models therefore forms an invaluable complement to epidemiology.

Golden Syrian hamsters are highly susceptible to infection with SARS-CoV-2, show clear signs of disease, shed the virus at high titers from the upper respiratory tract, and transmit SARS-CoV-2 to direct contacts and through the air (18-20). This model species has been used to evaluate the utility of vaccines, drugs, and passively transferred antibodies for blocking transmission (20-22). Nevertheless, an important limitation of the model is that its extreme permissiveness can interfere with the detection of differences in transmission efficiency. To date, analysis of the fitness of novel variants has relied on their co-infection with ancestral strains (23-25). While sensitive, this approach introduces the likelihood of viral fitness being modulated by interactions between the co-infecting viruses (26, 27). There is therefore a need for refinement of the hamster model to increase the stringency of transmission. Identification of temporal and environmental factors that modulate transmission can help to achieve this goal.

Here, we used a hamster model to investigate the impact of a range of temporal and environmental conditions on SARS-CoV-2 transmission. Our data indicate that high transmission efficiency is maintained with exposure durations as short as one hour and across a wide range of humidity and temperature conditions. Conversely, the timing of exposure relative to infection of donor animals was found to strongly modulate transmission, with exposure prior to 16 h and after 48 hours post-inoculation (hpi) yielding little or no spread to contacts. The temporal structure of transmission corresponded with infectious viral titers in the nasal tract, strongly suggesting that the infectious period is defined by the dynamics of viral load in donor hosts.

## Results

### Minimal impact of exposure duration on SARS-CoV-2 transmission

Reasoning that viral densities in the air or the recipient respiratory tract may accumulate over time, we hypothesized that the period of time during which naïve animals are exposed to the exhaled breath of infected animals would impact the efficiency of transmission. To test this hypothesis, we used an exposure system in which naïve hamsters are placed on the opposite side of a porous, double-walled, barrier from infected hamsters for a defined duration (**Supplementary Figure 1**). Exposures were carried out under controlled environmental conditions at 24 hpi of donor animals for periods of 5 d, 8 h, 4 h and 1 h. After exposure, animals were singly housed. Collection of nasal lavage samples was used to monitor for transmission. The results revealed robust transmission for all exposure durations tested (**Figure 1**). Positivity in a subset of contact animals was detected at the first sampling time of 2 d post-inoculation (1 d post-exposure) and all contact animals were found to harbor infectious virus by 8 d post-inoculation (7 d post-exposure). The period of exposure did not give rise to clear temporal trends in transmission. Thus, within the range tested, the period of exposure had minimal impact on SARS-CoV-2 transmission in hamsters. The data indicate that a minimal infectious dose is readily transferred to recipient hamsters within the course of one hour.

**Figure 1.**
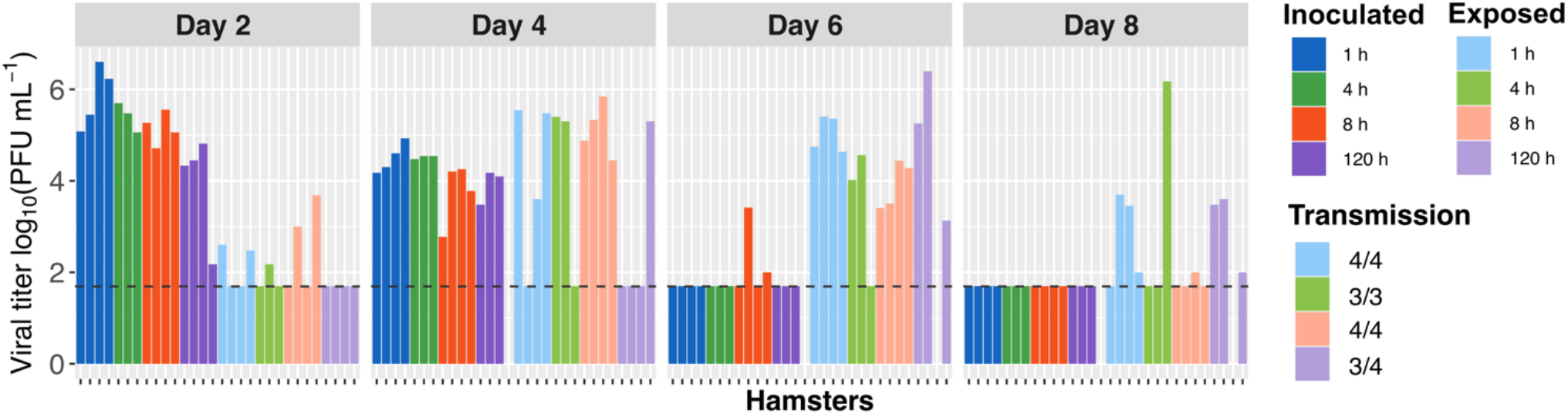
Exposure durations as short as 1 h are sufficient for robust SARS-CoV-2 transmission. Viral titers in nasal lavage samples collected from inoculated (dark colors) and exposed (light colors) hamsters are plotted. Facets show results from 2, 4, 6 and 8 days post-inoculation (dpi) with different colors indicating the duration of exposure. Animals were inoculated with 1×10^4^ PFU (titered on VeroE6 cells) and exposures initiated at 24 hpi. Exposures were carried out for five days at 30°C and 50% RH. Horizontal dashed line indicates limit of detection (50 PFU). Missing data indicate that the animal died or was euthanized mid-way through the experiment. The fraction of transmission pairs in which recipients shed infectious virus at one or more time points is indicated at the right.

### Timing of exposure strongly modulates SARS-CoV-2 transmission

We next tested the impact of the timing of exposure on SARS-CoV-2 transmission. To test for transmission at early times, brief exposures of one or two hours were carried out at 10–12, 12– 14, 14–16, 16–17 or 24–25 hpi. Similarly, to evaluate transmission potential late in the course of infection, exposures were carried out for two hours on days 2, 4 and 6 post-inoculation. Nasal lavage samples collected from donor animals at the conclusion of each exposure window were used to assess viral loads at the time of exposure, while serial samples collected from these same donors across multiple time points were used to evaluate the dynamics of viral load.

Analysis of nasal viral load in donor animals sampled at the time of exposure revealed a sharp increase between 12 h and 25 hpi, from an average of less than 1×10^3^ PFU/ml at 12 h to approximately 1×10^7^ PFU/ml at 25 h. Loads then declined over 2, 4 and 6 days post-inoculation (dpi), averaging about 1×10^6^ PFU/ml on day 2 and declining to < 1×10^2^ PFU/ml by day 6 (**Figure 2A**). Dynamics of viral load across all inoculated animals sampled on days 2, 4, 6 and 8 were consistent with these results obtained at the conclusion of specific exposures and revealed that titers were typically below the limit of detection by 8 dpi (**Supplementary Figure 2**). Taken together, the results show high viral loads of 1×10^5^ – 1×10^7^ PFU/ml are reached by 17 h and sustained at 48 hpi, but that titers fall below this range by 4 dpi.

**Figure 2.**
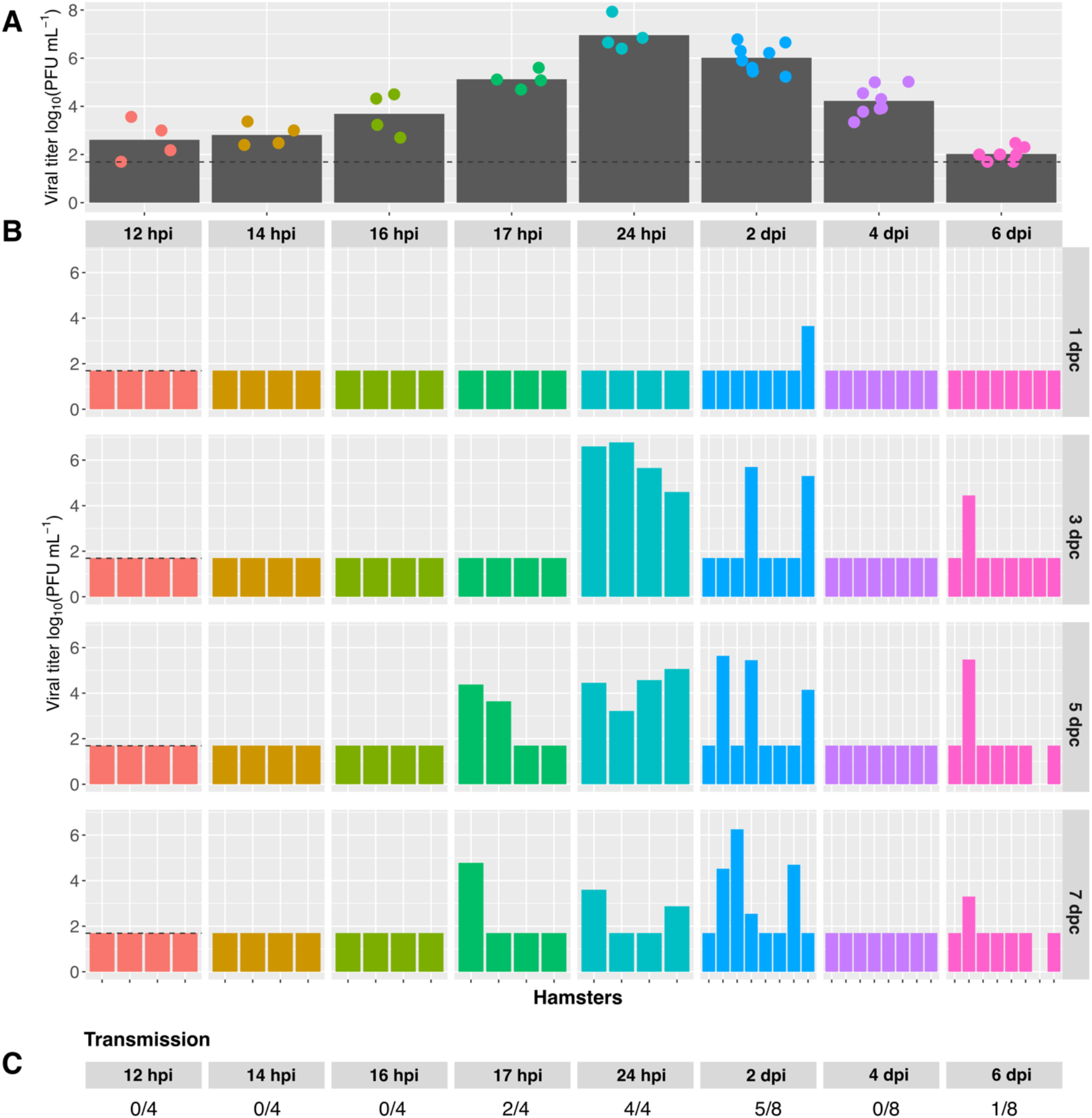
Period of transmissibility corresponds to period of high viral load in donor animals. A) Viral load in inoculated animals at the conclusion of the exposure period. Time points are indicated under the x-axis and a different color is assigned to each different exposure period. Bars show mean viral titer and dots show individual hamsters. B) Viral titers in nasal washes collected from contacts. Different exposure periods are shown with different colors. The time post-inoculation at which the exposure period concluded is shown at the top of each facet. Time points at which nasal wash samples were collected are shown at the right of each row in units of days post contact (dpc). For the early exposure groups, n=4 transmission pairs. For the 2 dpi, 4 dpi and 6 dpi exposure groups, data from two independent experiments are displayed together giving total n=8 transmission pairs. All animals were inoculated with 1×10^2^ PFU (titered on VeroE6 cells) and exposures were carried out at 20°C and 50% RH. Horizontal dashed line indicates limit of detection (50 PFU). Missing data indicate that the animal died or was euthanized mid-way through the experiment. C) The fraction of transmission pairs in which recipients shed infectious virus at one or more time points is indicated.

Analysis of exposed animals revealed a strong impact of the timing of exposure on transmission (**Figure 2B**). Naïve animals exposed for 2 h beginning at 10, 12 or 14 hpi of donors were not infected. By contrast, transmission was seen in two of four animals exposed for 1 h beginning at 16 hpi and all four animals exposed for 1 h beginning at 24 hpi. Exposure at late time points showed appreciable transmission only on day 2 post-inoculation, with five of eight hamsters contracting infection when exposed for 2 h beginning at 2 dpi. Overall, the data reveal a window of transmissibility from 17 h to 2 d after infection of the donor hamster (**Figure 2C**).

To test whether viral load is a likely determinant of infectious period, nasal lavage titers in donor animals at the conclusion of exposure were evaluated. Significantly higher viral loads were detected in hamsters that transmitted to contacts, compared to those that did not transmit SARS-CoV-2 (**Figure 3**).

**Figure 3.**
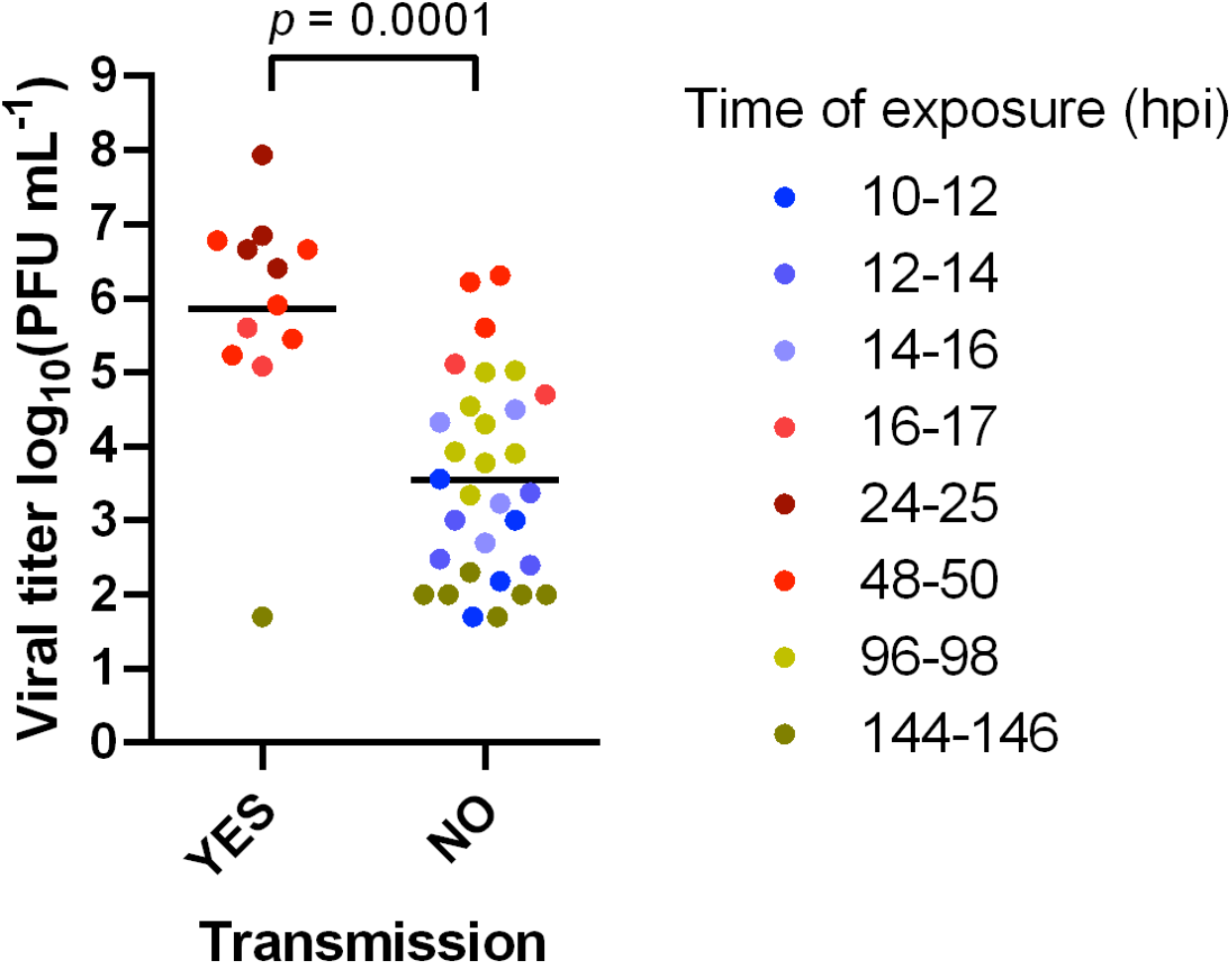
Transmission is associated with higher donor viral loads. Data analyzed are also shown in Figure 2. Viral titers detected in nasal lavage samples of donor animals collected at the time of exposure are plotted according to whether transmission occurred or not. All animals were inoculated with 1×10^2^ PFU (titered on VeroE6 cells) and exposures were carried out at 20°C and 50% RH. The time of exposure in hours post-inoculation (hpi) is shown in different colors. Unpaired Student’s t-test was performed on log-transformed data.

### Minimal impact of humidity on SARS-CoV-2 transmission

Ambient humidity is known to modulate the efficiency of aerosol transmission of influenza A virus, with dry conditions favoring spread (28-30). We hypothesized that a similar effect would be detectable for SARS-CoV-2 and therefore evaluated transmission with exposures carried out under dry (20 or 30% RH), intermediate (50% RH) or humid (80% RH) conditions. In each case, temperature was held constant at 20°C. When a five day exposure duration beginning at 24 hpi was used, transmission was found to be highly efficient irrespective of humidity (**Figure 4A**). Reasoning that any effects of humidity may be difficult to detect when overall transmission efficiencies are saturated, we performed similar experiments with exposures carried out early after infection when intermediate transmission frequency was observed. Namely, naïve animals were exposed for three hours beginning at 14 h after inoculation of donors. Since placement of the hamsters disrupts the relative humidity in their environment, this timing allowed equilibration of relative humidity during a period when no transmission was observed (14–16 hpi; **Figure 2B**). Here, results revealed a trend of more transmission under high RH conditions (**Figure 4B**). These results clearly indicated that, contrary to expectation, low ambient humidity was not favorable for transmission. However, hamsters were subjected to humidity set points for only a brief period, during exposure. If the effects of humidity on transmission act at the level of the host, then they would be unlikely to be manifested in this experimental system. To address this possibility, we tested whether pre-conditioning of animals under different humidity conditions for 4 d prior to inoculation or exposure impacted transmission. For a given group of hamsters, pre-conditioning and exposure were carried out under the same dry, intermediate or humid conditions. Again, the most transmission was seen at high RH (**Figure 4C**). These results suggest that high ambient RH may support SARS-CoV-2 transmission in this model and indicate that low RH does not promote transmission in this system.

**Figure 4.**
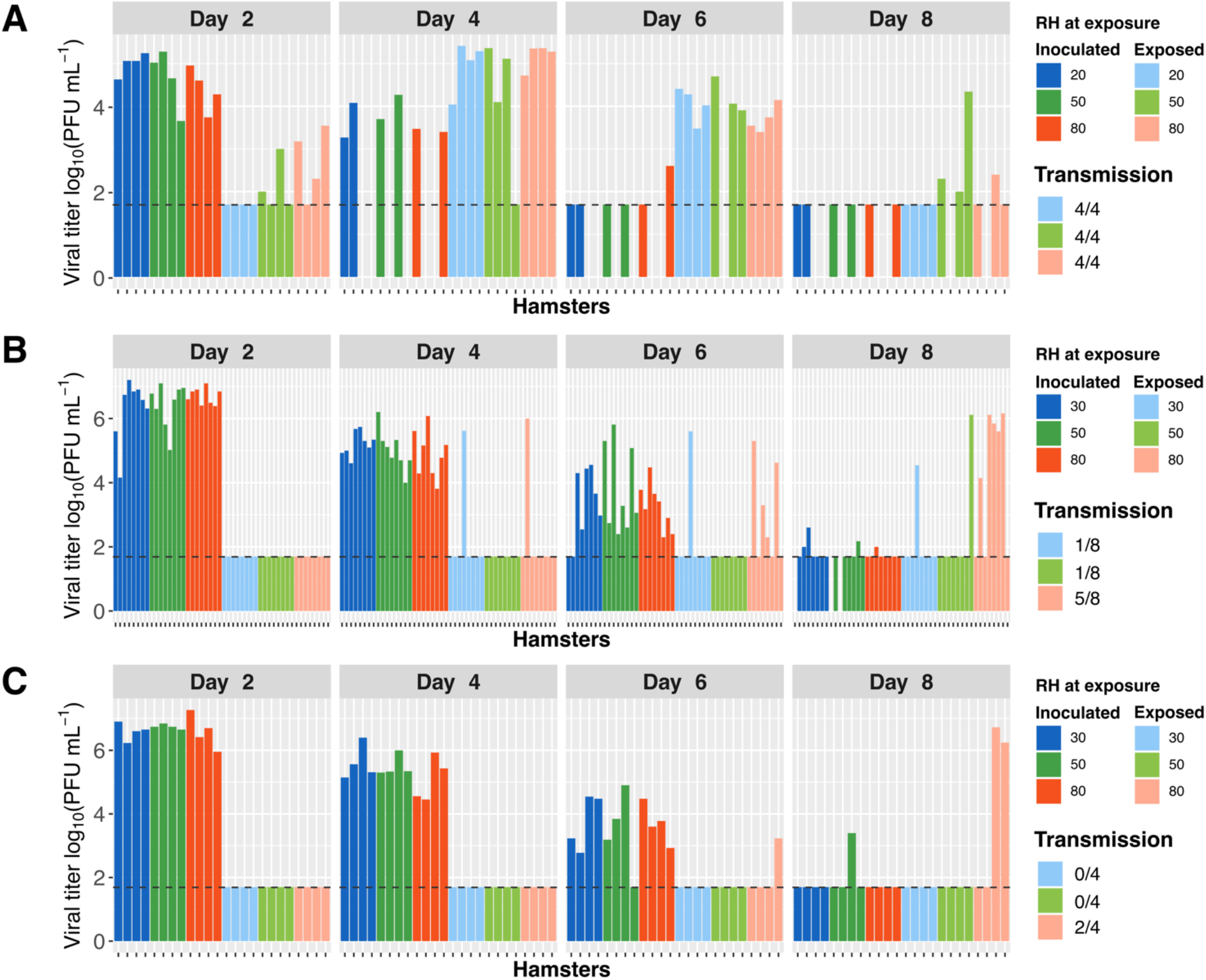
Low humidity does not promote SARS-CoV-2 transmission in hamsters. Viral titers in nasal lavage samples collected from inoculated (dark colors) and exposed (light colors) hamsters are plotted. Facets show results from 2, 4, 6 and 8 days post-inoculation (dpi) with different colors indicating the RH of exposure. A) Donor animals were inoculated with 1×10^4^ PFU (titered on VeroE6 cells) and contacts were exposed for a period five days under the indicated RH conditions, beginning at 24 hpi. N=4 transmission pairs. B) Donor animals were inoculated with 1×10^2^ PFU (titered on VeroE6 cells) and contacts were exposed for a period of 3 h under the indicated RH conditions, beginning at 14 hpi. Data are combined from two independent experiments, giving a total of n=8 transmission pairs. C) Donor and contact hamsters were preconditioned to the tested environmental RH for a period of four days. Donor animals were then inoculated with 1×10^2^ PFU (titered on VeroE6 cells) and contacts were exposed for a period 3 h under the indicated RH conditions, beginning at 14 hpi. N=4 transmission pairs. Horizontal dashed line indicates limit of detection (50 PFU). Missing data indicate that the animal died or was euthanized mid-way through the experiment. Temperature was 20°C in all experiments shown.

### Minimal impact of temperature on SARS-CoV-2 transmission

As for RH, ambient temperature was previously found to modulate the efficiency of aerosol transmission of influenza A virus, with cold conditions favoring spread (28-30). We hypothesized that a similar effect would be detectable for SARS-CoV-2 and therefore evaluated transmission with exposures carried out under cold (5°C), intermediate (20°C) or warm (30°C) conditions. In each case, RH was held constant at 50%. When a five-day exposure duration beginning at 24 hpi was used to compare 20°C and 30°C environments, transmission was found to be highly efficient under both conditions (**Figure 5A**). Since our hypothesis was that cold conditions would be favorable, we did not test 5°C in this system. Instead, we performed similar experiments with a one-hour exposure beginning at 16 hpi – an approach designed to yield intermediate levels of transmission and thereby allow detection of temperature effects. While intermediate levels of transmission were observed at 20°C and 30°C in these experiments, no transmission was observed at 5°C. Thus, results did not support the hypothesis that cold conditions augment transmission of SARS-CoV-2 (**Figure 5B**).

**Figure 5.**
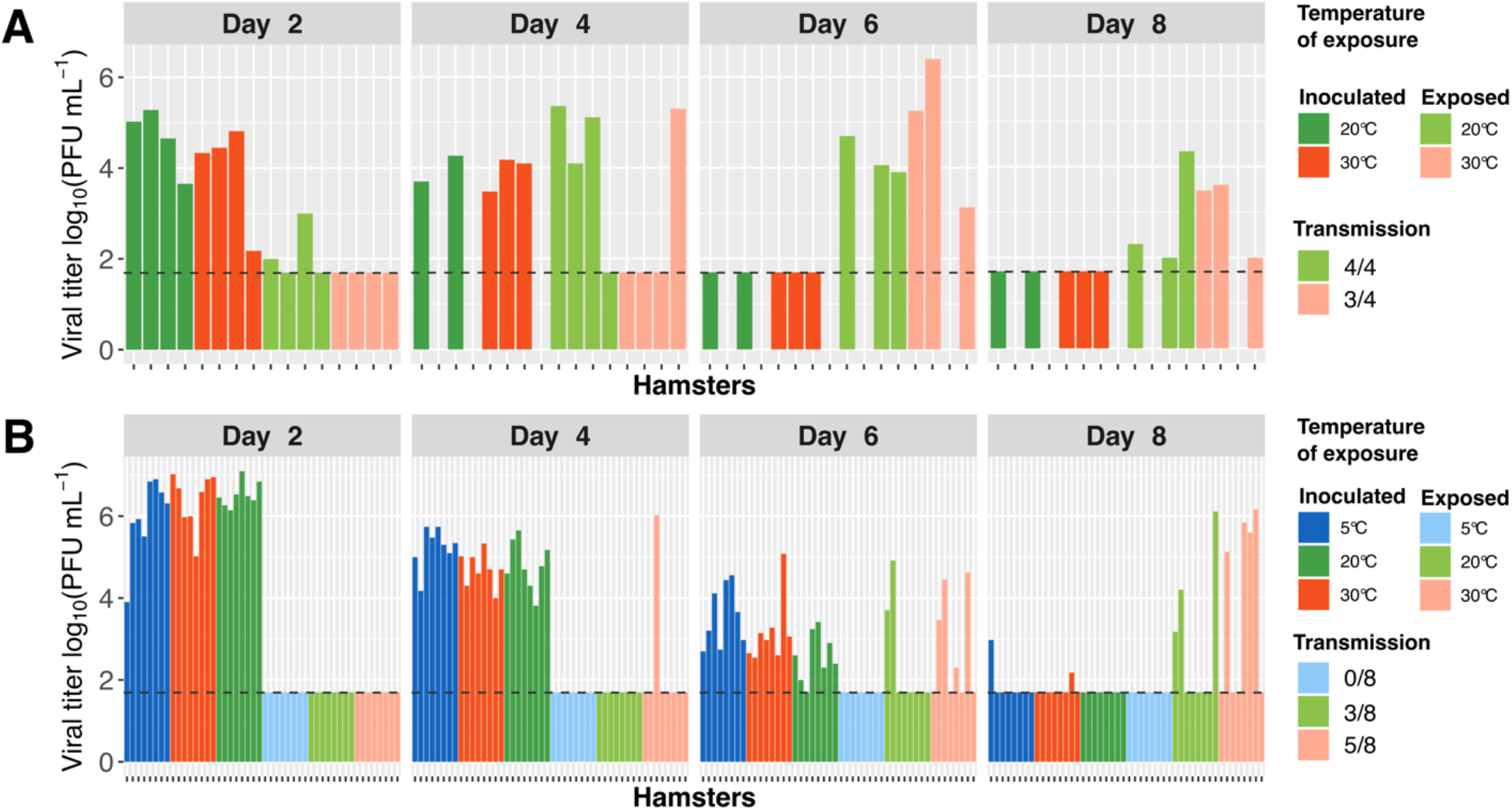
Low temperature does not promote SARS-CoV-2 transmission in hamsters. Viral titers in nasal lavage samples collected from inoculated (dark colors) and exposed (light colors) hamsters are plotted. Facets show results from 2, 4, 6 and 8 days post-inoculation (dpi) with different colors indicating the temperature of exposure. A) Donor hamsters were inoculated with 1×10^4^ PFU (titered on VeroE6 cells) and contacts were exposed for a period five days under the indicated temperature conditions, beginning at 24 hpi. N=4 transmission pairs. B) Donor animals were inoculated with 1×10^2^ PFU (titered on VeroE6 cells) and contacts were exposed for a period of one hour under the indicated temperature conditions, from 16-17 hpi. Data are combined from two independent experiments, giving a total of n=8 transmission pairs. Horizontal dashed line indicates limit of detection (50 PFU). Missing data indicate that the animal died or was euthanized mid-way through the experiment. RH was 50% for all experiments shown.

## Discussion

The COVID-19 pandemic has laid bare the importance of understanding the efficiency and dynamics of respiratory virus transmission. Here we report the results of detailed animal studies designed to quantify the effects of exposure duration, exposure timing and environmental conditions on SARS-CoV-2 transmission. Using a hamster model in which donor and contact animals share air space but are not in direct contact, we find that SARS-CoV-2 transmission is highly efficient, leading to infection of the majority of recipient animals with exposure periods as short as one hour and under a wide range of humidity and temperature conditions. Peak transmission was observed when exposures were carried out between 16 h and 48 h after inoculation of donor hamsters, revealing an early and narrow window of opportunity for transmission. This period of infectivity corresponded with high viral titers in the nasal tract, implicating viral load as a major driver of transmission.

Substantial evidence has accumulated for pre-symptomatic transmission of SARS-CoV-2 among humans, suggesting that the infectious period begins early after infection (31-34). However, the timing of infection and any subsequent transmission are often difficult to determine in natural settings. Experimental studies allow these parameters to be determined with precision and our data indicate that onward transmission of SARS-CoV-2 occurs readily after only a brief incubation period of ∼16 h. However, in translating this information to humans, it is important to note that the course of viral replication observed in inoculated hamsters was abbreviated relative to that in humans. While clinical data suggest that peak viral loads occur 3–4 days after infection (35, 36), peak titers were seen in hamsters at 24 hpi. This difference of kinetics may relate to initial dose, as inoculation of 1×10^2^–1×10^4^ PFU used here likely exceed a typical natural dose. Indeed, examination of viral titers in recipient animals shows delayed kinetics of replication for brief, early exposures – likely associated with lower initial dose – compared to exposures carried out over an extended period or at the peak of donor viral load. As a result, the period of contagiousness identified in hamsters may not translate directly to human hosts. However, the observation that the period of contagiousness corresponds to times of high viral load is likely to extend to humans.

Our data suggest that temporal changes in the potential for transmission likely stem from changes in viral load. While a direct relationship between viral load and transmission potential is intuitive, the extent to which respiratory virus transmission relies on symptoms and is limited by antiviral responses in the donor individual remain active areas of investigation (37-40). The observation herein that transmission efficiency declines with infectious viral titers both early and late in the course of infection suggests that viral load is a primary determinant of transmission, irrespective of the host processes that influence viral load. These results are consistent with those of a COVID-19 cohort study which showed a strong relationship between transmission and viral load, independent of symptoms (41). Viral loads detected in infected individuals vary across several orders of magnitude (36, 42); while timing and method of sample collection likely contribute to this disparity, biological variation appears to be high. The link between viral loads and transmission is therefore critical to understand. Our data support the notion that heterogeneity in viral load across individuals is a plausible driver of the extreme transmission heterogeneity observed for SARS-CoV-2 (31, 43).

Similarities between coronaviruses and influenza viruses in their modes of transmission, particle structure and seasonality suggest that similar factors likely shape the transmission of these pathogens (44). For influenza A virus (IAV), we previously showed that ambient temperature and humidity have a strong impact on transmission in controlled, experimental settings (28-30). Clear correlations between these meteorological variables and population level influenza activity have also been reported (45, 46). Thus, seasonal changes in humidity and temperature are likely major drivers of influenza dynamics at the population level. These observations led to our hypothesis that SARS-CoV-2 transmission would be suppressed by high ambient humidity and temperature conditions. Our data do not validate this hypothesis and instead reveal efficient transmission among hamsters housed in humid or warm environments. Robust transmission under humid conditions is consistent with the U-shaped relationship between RH and stability that has been reported for multiple enveloped viruses, including SARS-CoV-2 (47-49). In general, however, these results were unexpected based on reports of super-spreading events in cold or dry indoor environments (50-53) and available information on how temperature and RH modulate viral stability, host susceptibility and aerosol dynamics (49, 54-56). The unexpected results may stem from our approach of performing exposures early in the course of infection; transmission success may be more subject to stochastic effects at early times compared to exposures carried out at times when viral load in donors is at peak levels.

While circulation of endemic coronaviruses shows a clear seasonal pattern (57-59), increased activity in winter has not been a consistent feature of the COVID-19 pandemic. Influenza pandemics also do not follow the typical seasonality of epidemic influenza but have shown a decline in circulation during summer months (60, 61), suggesting the relative influence of seasonal drivers is reduced but not ablated for pandemic influenza. In the case of SARS-CoV-2, a lack of seasonal patterns at this early stage in its circulation may reflect low population immunity, dominance of other epidemiological factors such as population behavior and government interventions (16), or a lack of sensitivity to the seasonal drivers that shape the dynamics of epidemic coronaviruses and seasonal influenza viruses. While our data suggest a lack of sensitivity to humidity and temperature, we caution against over-interpretation of these data given the limitations inherent in our experimental system.

A detailed understanding of the host, viral and environmental factors that shape transmission efficiency is of fundamental importance to efforts to elucidate the drivers of SARS-CoV-2 dynamics across spatial-temporal scales. This knowledge in turn is invaluable for refining strategies to interrupt transmission. Our data reveal that the timing of exposure is a potent determinant of transmission potential and point to viral load as the underlying driver of this effect.

## Methods

### Virus

SARS-CoV-2/human/USA/GA-83E/2020 was isolated from a clinical sample by culture on VeroE6 cells (62). The passage 1 virus stock was aliquoted and stored at −80°C. Genomic analysis verified the predominance of full-length genomes and retention of the furin cleavage site in Spike. Sequencing also revealed the presence of the D614G polymorphism in the Spike protein and six nucleotide deletion in ORF 8 (nucleotides 28,090-28,095; amino acids AGSKS to A--ES). Viral titers were determined by plaque assay on VeroE6 cells. The passage 1 virus stock was used for all experiments outlined herein.

### Cells

VeroE6 cells were obtained from ATCC (clone E6, ATCC, #CRL-1586) and cultured in DMEM (Gibco) supplemented with 10% fetal bovine serum (Biotechne), MEM nonessential amino acids (Corning) and Normocin (Invivogen). Cells were routinely confirmed to be negative for mycoplasma contamination. These cells were used for viral culture and titration.

### Animals

All animal experiments were conducted in accordance with the Guide for the Care and Use of Laboratory Animals of the National Institutes of Health. Studies were conducted under animal biosafety level 3 (ABSL-3) containment and approved by the IACUC of Emory University (protocol PROTO202000055). Animals were humanely euthanized following guidelines approved by the American Veterinary Medical Association (AVMA). Outbred, male, Golden Syrian hamsters of 90-101 g body weight were obtained from Charles River Laboratories and singly housed on paper bedding with access to food and water *ad libitum*. Prior to inoculation, nasal lavage and euthanasia, hamsters were sedated with ketamine (120 mg/kg) – xylazine (4 mg/kg) administered intraperitoneally. Xylazine was reversed with 1mg/kg of atipamezole administered intraperitoneally. Animal health was monitored daily through visual observation and determination of body weight.

### Inoculation

Virus was diluted serially in PBS to achieve the desired dose and then sedated hamsters were inoculated intranasally with a 100 ul, applied dropwise to both nares with the animal in dorsal recumbency. Doses ranged from 1×10^2^ PFU to 1×10^4^ PFU (titered on VeroE6 cells), as indicated in figure legends.

### Nasal lavage

Virus was sampled from the upper respiratory tract by nasal lavage. Sedated hamsters were placed in ventral recumbency with nose suspended above an open Petri dish. A total volume of 400 ul PBS was applied to the nares using a micropipette and allowed to drop back into the dish. An additional volume of 200 ul PBS was used to wash the dish. Fluid in the dish was collected, aliquoted and stored at −80°C prior to determination of viral titers by plaque assay.

### Exposure of naïve hamsters to inoculated hamsters

Exposures were carried out in rodent cages modified through the addition of a double-walled porous barrier, which divided the cage in two. A single hamster was placed on either side of the barrier. The barrier reached from wall-to-wall and floor-to-lid and comprised two stainless-steel sheets placed 1 cm apart, with 0.5 cm perforations arrayed across each sheet. Each side of the cage was supplied with food and water. The cage was enclosed with a filter top.

For all exposures, cages were placed within a Caron 6040 environmental chamber. These chambers allow tight control of humidity and temperature conditions. To achieve uniformity of these conditions throughout the chamber, air flow rates are relatively high, at 13000 L/min. Environmental conditions within the chamber and within the rodent cages (with hamsters present) were verified using a Temperature/Humidity WIFI Data Logger (Traceable Products; Webster, Texas). Desired conditions were readily met within the hamster cages. After opening the chamber door to place cages, recovery time varied from 10 to 60 minutes, with dry (20 or 30% RH) conditions requiring the longest recovery times. To allow testing of different RH conditions, it was therefore important to place the animals in the chambers at an early time point after inoculation of donors (14 hpi) such that chamber equilibration was achieved prior to the start of the infectious period (16 hpi).

Inoculated donor animals were singly housed within environmental chambers shortly after inoculation. One naïve recipient was introduced on the opposite side of each exposure cage at 14 h – 6 dpi, as indicated in each figure legend. Exposures were carried out for durations ranging from 1 h to 5 d. Where exposures exceeded 24 h, animals were removed from the chambers on a daily basis for determination of body weight and, on a subset of days, nasal lavage. To avoid spurious transmission, care was taken during animal handling. Gloves were changed and biosafety cabinet and weighing container were disinfected between animals.

## Data analysis

Data analysis was done using RStudio 1.3.959 and Prism version 6.0.7. Plots were aesthetically modified using Inkscape 1.0.

## Acknowledgements

We thank Hui Tao and Shamika Danzy for technical assistance and the Emory University Division of Animal Resources for their support with animal care, cage construction and logistical challenges. This work was funded by the National Institute of Allergy and Infectious Disease through the Centers of Excellence for Influenza Research and Surveillance (CEIRS) contract no. HHSN272201400004C. GKD is supported by F31 AI 50114.

## References

1. Flaxman S, Mishra S, Gandy A, Unwin HJT, Mellan TA, Coupland H, Whittaker C, Zhu H, Berah T, Eaton JW, Monod M, Perez-Guzman PN, Schmit N, Cilloni L, Ainslie KEC, Baguelin M, Boonyasiri A, Boyd O, Cattarino L, Cooper LV, Cucunubá Z, Cuomo-Dannenburg G, Dighe A, Djaafara B, Dorigatti I, van Elsland SL, FitzJohn RG, Gaythorpe KAM, Geidelberg L, Grassly NC, Green WD, Hallett T, Hamlet A, Hinsley W, Jeffrey B, Knock E, Laydon DJ, Nedjati-Gilani G, Nouvellet P, Parag KV, Siveroni I, Thompson HA, Verity R, Volz E, Walters CE, Wang H, Wang Y, Watson OJ, Winskill P, Xi X, et al. 2020. Estimating the effects of non-pharmaceutical interventions on COVID-19 in Europe. Nature 584:257–261.

2. Lai S, Ruktanonchai NW, Zhou L, Prosper O, Luo W, Floyd JR, Wesolowski A, Santillana M, Zhang C, Du X, Yu H, Tatem AJ. 2020. Effect of non-pharmaceutical interventions to contain COVID-19 in China. Nature 585:410–413.

3. Ferguson N, Laydon D, Nedjati Gilani G, Imai N, Ainslie K, Baguelin M, Bhatia S, Boonyasiri A, Cucunuba Perez Z, Cuomo-Dannenburg G. 2020. Report 9: Impact of non-pharmaceutical interventions (NPIs) to reduce COVID19 mortality and healthcare demand.

4. Ng Y, Li Z, Chua YX, Chaw WL, Zhao Z, Er B, Pung R, Chiew CJ, Lye DC, Heng D, Lee VJ. 2020. Evaluation of the Effectiveness of Surveillance and Containment Measures for the First 100 Patients with COVID-19 in Singapore - January 2-February 29, 2020. MMWR Morb Mortal Wkly Rep 69:307–311.

5. Ng OT, Marimuthu K, Koh V, Pang J, Linn KZ, Sun J, De Wang L, Chia WN, Tiu C, Chan M, Ling LM, Vasoo S, Abdad MY, Chia PY, Lee TH, Lin RJ, Sadarangani SP, Chen MI, Said Z, Kurupatham L, Pung R, Wang LF, Cook AR, Leo YS, Lee VJ. 2021. SARS-CoV-2 seroprevalence and transmission risk factors among high-risk close contacts: a retrospective cohort study. Lancet Infect Dis 21:333–343.

6. Quilty BJ, Clifford S, Hellewell J, Russell TW, Kucharski AJ, Flasche S, Edmunds WJ, Centre for the Mathematical Modelling of Infectious Diseases C-wg. 2021. Quarantine and testing strategies in contact tracing for SARS-CoV-2: a modelling study. Lancet Public Health 6:e175–e183.

7. Ashcroft P, Lehtinen S, Angst DC, Low N, Bonhoeffer S. 2021. Quantifying the impact of quarantine duration on COVID-19 transmission. Elife 10.

8. Ma Y, Pei S, Shaman J, Dubrow R, Chen K. 2021. Role of meteorological factors in the transmission of SARS-CoV-2 in the United States. Nature Communications 12:3602.

9. Choi Y-W, Tuel A, Eltahir EAB. 2021. On the Environmental Determinants of COVID-19 Seasonality. GeoHealth 5:e2021GH000413–e2021GH000413.

10. Smith TP, Flaxman S, Gallinat AS, Kinosian SP, Stemkovski M, Unwin HJT, Watson OJ, Whittaker C, Cattarino L, Dorigatti I, Tristem M, Pearse WD. 2021. Temperature and population density influence SARS-CoV-2 transmission in the absence of nonpharmaceutical interventions. Proceedings of the National Academy of Sciences 118:e2019284118.

11. Smith TP, Flaxman S, Gallinat AS, Kinosian SP, Stemkovski M, Unwin HJT, Watson OJ, Whittaker C, Cattarino L, Dorigatti I, Tristem M, Pearse WD. 2021. Temperature and population density influence SARS-CoV-2 transmission in the absence of nonpharmaceutical interventions. Proc Natl Acad Sci U S A 118.

12. Landier J, Paireau J, Rebaudet S, Legendre E, Lehot L, Fontanet A, Cauchemez S, Gaudart J. 2021. Cold and dry winter conditions are associated with greater SARS-CoV-2 transmission at regional level in western countries during the first epidemic wave. Sci Rep 11:12756.

13. Yao Y, Pan J, Liu Z, Meng X, Wang W, Kan H, Wang W. 2020. No association of COVID-19 transmission with temperature or UV radiation in Chinese cities. Eur Respir J 55.

14. Juni P, Rothenbuhler M, Bobos P, Thorpe KE, da Costa BR, Fisman DN, Slutsky AS, Gesink D. 2020. Impact of climate and public health interventions on the COVID-19 pandemic: a prospective cohort study. CMAJ 192:E566–E573.

15. Yuan J, Wu Y, Jing W, Liu J, Du M, Wang Y, Liu M. 2021. Association between meteorological factors and daily new cases of COVID-19 in 188 countries: A time series analysis. Sci Total Environ 780:146538.

16. Sera F, Armstrong B, Abbott S, Meakin S, O’Reilly K, von Borries R, Schneider R, Roye D, Hashizume M, Pascal M, Tobias A, Vicedo-Cabrera AM, Network MCCCR, Group CC-W, Gasparrini A, Lowe R. 2021. A cross-sectional analysis of meteorological factors and SARS-CoV-2 transmission in 409 cities across 26 countries. Nat Commun 12:5968.

17. Mecenas P, Bastos R, Vallinoto ACR, Normando D. 2020. Effects of temperature and humidity on the spread of COVID-19: A systematic review. PLoS One 15:e0238339.

18. Sia SF, Yan L-M, Chin AW, Fung K, Poon LL, Nicholls JM, Peiris M, Yen H-L. Pre-print. Pathogenesis and transmission of SARS-CoV-2 virus in golden Syrian hamsters. doi:10.21203/rs.3.rs-20774/v1.

19. Imai M, Iwatsuki-Horimoto K, Hatta M, Loeber S, Halfmann PJ, Nakajima N, Watanabe T, Ujie M, Takahashi K, Ito M, Yamada S, Fan S, Chiba S, Kuroda M, Guan L, Takada K, Armbrust T, Balogh A, Furusawa Y, Okuda M, Ueki H, Yasuhara A, Sakai-Tagawa Y, Lopes TJS, Kiso M, Yamayoshi S, Kinoshita N, Ohmagari N, Hattori SI, Takeda M, Mitsuya H, Krammer F, Suzuki T, Kawaoka Y. 2020. Syrian hamsters as a small animal model for SARS-CoV-2 infection and countermeasure development. Proc Natl Acad Sci U S A 117:16587–16595.

20. Lee CY, Lowen AC. 2021. Animal models for SARS-CoV-2. Curr Opin Virol 48:73–81.

21. Horiuchi S, Oishi K, Carrau L, Frere J, Moller R, Panis M, tenOever BR. 2021. Immune memory from SARS-CoV-2 infection in hamsters provides variant-independent protection but still allows virus transmission. Sci Immunol doi:10.1126/sciimmunol.abm3131:eabm3131.

22. Abdelnabi R, Foo CS, Kaptein SJF, Zhang X, Do TND, Langendries L, Vangeel L, Breuer J, Pang J, Williams R, Vergote V, Heylen E, Leyssen P, Dallmeier K, Coelmont L, Chatterjee AK, Mols R, Augustijns P, De Jonghe S, Jochmans D, Weynand B, Neyts J. 2021. The combined treatment of Molnupiravir and Favipiravir results in a potentiation of antiviral efficacy in a SARS-CoV-2 hamster infection model. EBioMedicine 72:103595.

23. Zhou B, Thao TTN, Hoffmann D, Taddeo A, Ebert N, Labroussaa F, Pohlmann A, King J, Steiner S, Kelly JN, Portmann J, Halwe NJ, Ulrich L, Trueb BS, Fan X, Hoffmann B, Wang L, Thomann L, Lin X, Stalder H, Pozzi B, de Brot S, Jiang N, Cui D, Hossain J, Wilson MM, Keller MW, Stark TJ, Barnes JR, Dijkman R, Jores J, Benarafa C, Wentworth DE, Thiel V, Beer M. 2021. SARS-CoV-2 spike D614G change enhances replication and transmission. Nature 592:122–127.

24. Hou YJ, Chiba S, Halfmann P, Ehre C, Kuroda M, Dinnon KH, 3rd, Leist SR, Schafer A, Nakajima N, Takahashi K, Lee RE, Mascenik TM, Graham R, Edwards CE, Tse LV, Okuda K, Markmann AJ, Bartelt L, de Silva A, Margolis DM, Boucher RC, Randell SH, Suzuki T, Gralinski LE, Kawaoka Y, Baric RS. 2020. SARS-CoV-2 D614G variant exhibits efficient replication ex vivo and transmission in vivo. Science 370:1464–1468.

25. Plante JA, Liu Y, Liu J, Xia H, Johnson BA, Lokugamage KG, Zhang X, Muruato AE, Zou J, Fontes-Garfias CR, Mirchandani D, Scharton D, Bilello JP, Ku Z, An Z, Kalveram B, Freiberg AN, Menachery VD, Xie X, Plante KS, Weaver SC, Shi PY. 2021. Spike mutation D614G alters SARS-CoV-2 fitness. Nature 592:116–121.

26. Valcarcel J, Ortin J. 1989. Phenotypic hiding: the carryover of mutations in RNA viruses as shown by detection of mar mutants in influenza virus. J Virol 63:4107–9.

27. DaPalma T, Doonan BP, Trager NM, Kasman LM. 2010. A systematic approach to virus-virus interactions. Virus Res 149:1–9.

28. Lowen AC, Mubareka S, Steel J, Palese P. 2007. Influenza virus transmission is dependent on relative humidity and temperature. PLoS Pathog 3:1470–6.

29. Lowen AC, Steel J, Mubareka S, Palese P. 2008. High temperature (30 degrees C) blocks aerosol but not contact transmission of influenza virus. J Virol 82:5650–2.

30. Steel J, Palese P, Lowen AC. 2011. Transmission of a 2009 pandemic influenza virus shows a sensitivity to temperature and humidity similar to that of an H3N2 seasonal strain. J Virol 85:1400–2.

31. Adam DC, Wu P, Wong JY, Lau EHY, Tsang TK, Cauchemez S, Leung GM, Cowling BJ. 2020. Clustering and superspreading potential of SARS-CoV-2 infections in Hong Kong. Nat Med 26:1714–1719.

32. Huang L, Zhang X, Zhang X, Wei Z, Zhang L, Xu J, Liang P, Xu Y, Zhang C, Xu A. 2020. Rapid asymptomatic transmission of COVID-19 during the incubation period demonstrating strong infectivity in a cluster of youngsters aged 16-23 years outside Wuhan and characteristics of young patients with COVID-19: A prospective contact-tracing study. J Infect 80:e1–e13.

33. Johansson MA, Quandelacy TM, Kada S, Prasad PV, Steele M, Brooks JT, Slayton RB, Biggerstaff M, Butler JC. 2021. SARS-CoV-2 Transmission From People Without COVID-19 Symptoms. JAMA Netw Open 4:e2035057.

34. Hart WS, Maini PK, Thompson RN. 2021. High infectiousness immediately before COVID-19 symptom onset highlights the importance of continued contact tracing. Elife 10.

35. Jones TC, Biele G, Muhlemann B, Veith T, Schneider J, Beheim-Schwarzbach J, Bleicker T, Tesch J, Schmidt ML, Sander LE, Kurth F, Menzel P, Schwarzer R, Zuchowski M, Hofmann J, Krumbholz A, Stein A, Edelmann A, Corman VM, Drosten C. 2021. Estimating infectiousness throughout SARS-CoV-2 infection course. Science 373.

36. Ke R, Martinez PP, Smith RL, Gibson LL, Mirza A, Conte M, Gallagher N, Luo CH, Jarrett J, Conte A, Liu T, Farjo M, Walden KKO, Rendon G, Fields CJ, Wang L, Fredrickson R, Edmonson DC, Baughman ME, Chiu KK, Choi H, Scardina KR, Bradley S, Gloss SL, Reinhart C, Yedetore J, Quicksall J, Owens AN, Broach J, Barton B, Lazar P, Heetderks WJ, Robinson ML, Mostafa HH, Manabe YC, Pekosz A, McManus DD, Brooke CB. 2021. Daily sampling of early SARS-CoV-2 infection reveals substantial heterogeneity in infectiousness. medRxiv doi:10.1101/2021.07.12.21260208.

37. Lau LL, Cowling BJ, Fang VJ, Chan KH, Lau EH, Lipsitch M, Cheng CK, Houck PM, Uyeki TM, Peiris JS, Leung GM. 2010. Viral shedding and clinical illness in naturally acquired influenza virus infections. J Infect Dis 201:1509–16.

38. Tsang TK, Fang VJ, Chan KH, Ip DK, Leung GM, Peiris JS, Cowling BJ, Cauchemez S. 2016. Individual Correlates of Infectivity of Influenza A Virus Infections in Households. PLoS One 11:e0154418.

39. Danzy S, Lowen AC, Steel J. 2021. A quantitative approach to assess influenza A virus fitness and transmission in guinea pigs. J Virol doi:10.1128/JVI.02320-20.

40. Harrison AG, Lin T, Wang P. 2020. Mechanisms of SARS-CoV-2 Transmission and Pathogenesis. Trends Immunol 41:1100–1115.

41. Marks M, Millat-Martinez P, Ouchi D, Roberts CH, Alemany A, Corbacho-Monne M, Ubals M, Tobias A, Tebe C, Ballana E, Bassat Q, Baro B, Vall-Mayans M C GB, Prat N, Ara J, Clotet B, Mitja O. 2021. Transmission of COVID-19 in 282 clusters in Catalonia, Spain: a cohort study. Lancet Infect Dis 21:629–636.

42. Jones TC, Biele G, Muhlemann B, Veith T, Schneider J, Beheim-Schwarzbach J, Bleicker T, Tesch J, Schmidt ML, Sander LE, Kurth F, Menzel P, Schwarzer R, Zuchowski M, Hofmann J, Krumbholz A, Stein A, Edelmann A, Corman VM, Drosten C. 2021. Estimating infectiousness throughout SARS-CoV-2 infection course. Science doi:10.1126/science.abi5273.

43. Lakdawala SS, Menachery VD. 2021. Catch me if you can: superspreading of COVID-19. Trends Microbiol doi:10.1016/j.tim.2021.05.002.

44. Moriyama M, Hugentobler WJ, Iwasaki A. 2020. Seasonality of Respiratory Viral Infections. Annu Rev Virol doi:10.1146/annurev-virology-012420-022445.

45. Shaman J, Goldstein E, Lipsitch M. 2011. Absolute humidity and pandemic versus epidemic influenza. Am J Epidemiol 173:127–35.

46. Shaman J, Pitzer VE, Viboud C, Grenfell BT, Lipsitch M. 2010. Absolute humidity and the seasonal onset of influenza in the continental United States. PLoS Biol 8:e1000316.

47. Prussin AJ, 2nd, Schwake DO, Lin K, Gallagher DL, Buttling L, Marr LC. 2018. Survival of the Enveloped Virus Phi6 in Droplets as a Function of Relative Humidity, Absolute Humidity, and Temperature. Appl Environ Microbiol 84.

48. Schaffer FL, Soergel ME, Straube DC. 1976. Survival of airborne influenza virus: effects of propagating host, relative humidity, and composition of spray fluids. Arch Virol 51:263–73.

49. Morris DH, Yinda KC, Gamble A, Rossine FW, Huang Q, Bushmaker T, Fischer RJ, Matson MJ, Van Doremalen N, Vikesland PJ, Marr LC, Munster VJ, Lloyd-Smith JO. 2021. Mechanistic theory predicts the effects of temperature and humidity on inactivation of SARS-CoV-2 and other enveloped viruses. Elife 10.

50. Dyal JW, Grant MP, Broadwater K, Bjork A, Waltenburg MA, Gibbins JD, Hale C, Silver M, Fischer M, Steinberg J, Basler CA, Jacobs JR, Kennedy ED, Tomasi S, Trout D, Hornsby-Myers J, Oussayef NL, Delaney LJ, Patel K, Shetty V, Kline KE, Schroeder B, Herlihy RK, House J, Jervis R, Clayton JL, Ortbahn D, Austin C, Berl E, Moore Z, Buss BF, Stover D, Westergaard R, Pray I, DeBolt M, Person A, Gabel J, Kittle TS, Hendren P, Rhea C, Holsinger C, Dunn J, Turabelidze G, Ahmed FS, deFijter S, Pedati CS, Rattay K, Smith EE, Luna-Pinto C, Cooley LA, et al. 2020. COVID-19 Among Workers in Meat and Poultry Processing Facilities - 19 States, April 2020. MMWR Morb Mortal Wkly Rep 69.

51. Gunther T, Czech-Sioli M, Indenbirken D, Robitaille A, Tenhaken P, Exner M, Ottinger M, Fischer N, Grundhoff A, Brinkmann MM. 2020. SARS-CoV-2 outbreak investigation in a German meat processing plant. EMBO Mol Med 12:e13296.

52. Atrubin D, Wiese M, Bohinc B. 2020. An Outbreak of COVID-19 Associated with a Recreational Hockey Game - Florida, June 2020. MMWR Morb Mortal Wkly Rep 69:1492–1493.

53. Khanh NC, Thai PQ, Quach HL, Thi NH, Dinh PC, Duong TN, Mai LTQ, Nghia ND, Tu TA, Quang N, Quang TD, Nguyen TT, Vogt F, Anh DD. 2020. Transmission of SARS-CoV 2 During Long-Haul Flight. Emerg Infect Dis 26:2617–2624.

54. Kormuth KA, Lin K, Prussin AJ, 2nd, Vejerano EP, Tiwari AJ, Cox SS, Myerburg MM, Lakdawala SS, Marr LC. 2018. Influenza Virus Infectivity Is Retained in Aerosols and Droplets Independent of Relative Humidity. J Infect Dis 218:739–747.

55. Kudo E, Song E, Yockey LJ, Rakib T, Wong PW, Homer RJ, Iwasaki A. 2019. Low ambient humidity impairs barrier function and innate resistance against influenza infection. Proc Natl Acad Sci U S A 116:10905–10910.

56. Marr LC, Tang JW, Van Mullekom J, Lakdawala SS. 2019. Mechanistic insights into the effect of humidity on airborne influenza virus survival, transmission and incidence. J R Soc Interface 16:20180298.

57. Li Y, Wang X, Nair H. 2020. Global Seasonality of Human Seasonal Coronaviruses: A Clue for Postpandemic Circulating Season of Severe Acute Respiratory Syndrome Coronavirus 2? The Journal of Infectious Diseases 222:1090–1097.

58. Monto AS, DeJonge PM, Callear AP, Bazzi LA, Capriola SB, Malosh RE, Martin ET, Petrie JG. 2020. Coronavirus Occurrence and Transmission Over 8 Years in the HIVE Cohort of Households in Michigan. The Journal of Infectious Diseases 222:9–16.

59. Li Y, Wang X, Nair H. 2020. Global Seasonality of Human Seasonal Coronaviruses: A Clue for Postpandemic Circulating Season of Severe Acute Respiratory Syndrome Coronavirus 2? J Infect Dis 222:1090–1097.

60. Fox SJ, Miller JC, Meyers LA. 2017. Seasonality in risk of pandemic influenza emergence. PLOS Computational Biology 13:e1005749.

61. Andreasen V, Viboud C, Simonsen L. 2008. Epidemiologic characterization of the 1918 influenza pandemic summer wave in Copenhagen: implications for pandemic control strategies. The Journal of infectious diseases 197:270–278.

62. Edara VV, Hudson WH, Xie X, Ahmed R, Suthar MS. 2021. Neutralizing Antibodies Against SARS-CoV-2 Variants After Infection and Vaccination. JAMA 325:1896–1898.

